# An analytical model describing the mechanics of erythrocyte membrane wrapping during active invasion of a plasmodium falciparum merozoite

**DOI:** 10.1101/2022.06.14.496094

**Authors:** Chimwemwe Msosa, Tamer Abdalrahman, Thomas Franz

## Abstract

The invasion of a merozoite into an erythrocyte by membrane wrapping is a hallmark of malaria pathogenesis. The invasion involves biomechanical interactions whereby the merozoite exerts actomyosin-based forces to push itself into and through the erythrocyte membrane while concurrently inducing biochemical damage to the erythrocyte membrane. Whereas the biochemical damage process has been investigated, the detailed mechanistic understanding of the invasion mechanics remains limited. Thus, the current study aimed to develop a mathematical model describing the mechanical factors involved in the merozoite invasion into an erythrocyte and explore the invasion mechanics.

A shell theory model was developed comprising constitutive, equilibrium and governing equations of the deformable erythrocyte membrane to predict membrane mechanics during the wrapping of an entire non-deformable ellipsoidal merozoite. Predicted parameters include principal erythrocyte membrane deformations and stresses, wrapping and indentation forces, and indentation work. The numerical investigations considered two limits for the erythrocyte membrane deformation during wrapping (4% and 51% areal strain) and erythrocyte membrane phosphorylation (decrease of membrane elastic modulus from 1 to 0.5 kPa).

For an intact erythrocyte, the maximum indentation force was 1 and 8.5 pN, and the indentation work was 1.92 ×10^-18^ and 1.40 ×10^-17^ J for 4% and 51% areal membrane strain. Phosphorylation damage in the erythrocyte membrane reduced the required indentation work by 50% to 0.97 ×10^-18^ and 0.70 ×10^-17^ J for 4% and 51% areal strain.

The current study demonstrated the developed model’s feasibility to provide new knowledge on the physical mechanisms of the merozoite invasion process that contribute to the invasion efficiency towards the discovery of new invasion-blocking anti-malaria drugs.

## 1 Introduction

The invasion of malaria merozoites into erythrocytes marks the pathogenesis of malaria. The invasion process involves biochemical and mechanical interactions to optimise the invasion efficiency. Biochemical interactions have been investigated extensively, whereas mechanical interactions between the host erythrocyte and the malaria merozoites received attention only more recently (Groomes et al., 2022). The deformation mechanics of erythrocytes membrane and how it affects the merozoite invasion process is virgin research point which need an intensive investigation. Merozoites have an actomyosin motor that promotes infiltration of erythrocytes by wrapping the deformable erythrocyte membrane with a deformable spectrin-actin layer to form a nascent parasitophorous vacuole (Hillringhaus et al., 2019; Koch and Baum, 2016; Weiss et al., 2015)

The invasion of a human erythrocyte by a malaria merozoite starts with a low-affinity contact mediated by the merozoite surface proteins (MSP) (Harvey et al., 2012). As soon as the merozoite is weakly attached to the erythrocyte surface, it begins to reorient to maximise the adhesion energy (Dasgupta et al., 2014). The adhesion energy is utilised to deform the erythrocyte membrane locally and, thus, form an invasion pit. Additionally, the reorientation of the merozoite facilitates the secretion of proteins located in specialised exocytic organelles of the malaria merozoite (micronemes, rhoptries) onto the apical surface of the erythrocyte (Singh et al., 2010). The molecular interaction between secreted erythrocyte binding antigen (EBA-175) and glycophorin antigen (GPA) (Jaskiewicz et al., 2019) plays a vital role in signalling the remodelling of the erythrocyte membrane skeleton (Zuccala et al., 2016). This is achieved by phosphorylation of the key membrane skeleton proteins such as ankyrin and spectrin.

Next, the tight junction, a circular structure that mechanically links the merozoite and erythrocyte membrane is established mediated by apical membrane antigen 1 (AMA1) binding to the rhoptry neck protein complex (RON2, 4, 5) (Richard et al., 2010; Srinivasan et al., 2011). Thereafter, the merozoite forms the digestive vacuole and exerts a traction force through the tight junction, causing the erythrocyte membrane to wrap around the merozoite surface. The tight junction moves rearward during the invasion, propelling the merozoite into the host cell (Cowman and Crabb, 2006). Predictions from imaging studies of merozoites fixed mid-entry indicate that an actomyosin force is applied consistently at or proximal to the tight junction (Riglar et al., 2011). In other words, the tight junction transmits the merozoite’s actomyosin force to the erythrocyte membrane, hence facilitating the invagination of the malaria merozoite (Koch and Baum, 2016).

Erythrocyte binding like ligands (EBL), erythrocyte binding antigens (EBA-175, EBA-140), and erythrocyte receptors such as glycophorin A (GPA) and glycophorin C (GPC) have been identified to mediate signalling and remodelling of host cytoskeletal components (Koch et al., 2017; Paul et al., 2015; Sisquella et al., 2017) whereas apical membrane antigen 1 (AMA1) and the rhoptry neck protein complex (RON2, 4, 5) are involved in formation of the tight junction (Richard et al., 2010; Srinivasan et al., 2011). Hence, these structures are vital in regulating invasion efficiency.

The internalisation of the malaria merozoite ends with the sealing of the erythrocyte membrane. The entire internalisation process of a merozoite into a human erythrocyte, illustrated schematically in Figure 1(a), takes less than 30 seconds.

**Figure 1:**
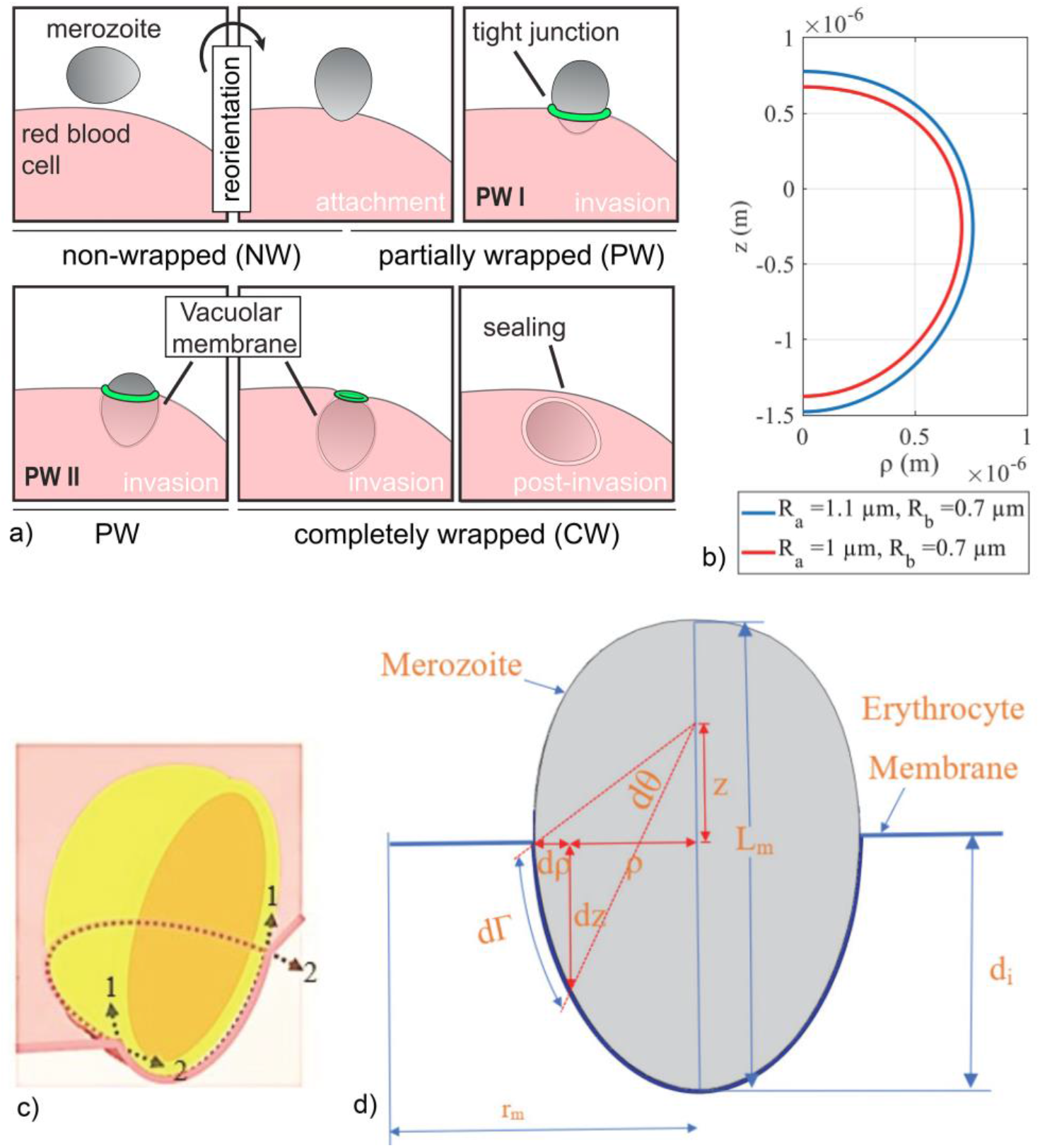
(a) Schematic illustration of invasion of malaria merozoite into a erythrocyte, from (Dasgupta et al., 2014). (b) Idealised merozoite shape with geometrical alteration resulting from altering R_a_ from 1 to 1.1 μm. (c) Principal stretch directions of the erythrocyte membrane, where 1 and 2 represent the meridian and circumferential directions, respectively. (d) An idealised schematic of merozoite partially wrapped by the erythrocyte membrane wrapping mechanism with L_m_ as total merozoite length and d_i_ as merozoite indentation depth.

The mechanistic research into erythrocyte invasion by merozoites is very scarce. An *in vitro* study by Li et al. (2014) into the invasion mechanics indicated that the yield point in quasi-statically loaded erythrocyte membranes ranged between 2% and 4% areal strain, beyond which lysis of the erythrocyte was observed.

Assessing the energetics of the internalisation of a malaria merozoite *in vitro* is challenging and costly. Mathematical modelling of the biomechanical interactions between the merozoite and erythrocyte can predict the force and energy required for invasion. Dasgupta et al. (2014) demonstrated that erythrocyte membrane wrapping is the primary mechanism for internalising the malaria merozoite. The authors suggested that the merozoite-induced remodelling of the erythrocyte cytoskeleton can use a considerable portion of the invasion energy. The total energy ψ required by the malaria merozoite to invade the erythrocyte comprises energies to overcome bending rigidity, membrane tension, adhesion strength, and line tension. The malaria merozoite must minimise the invasion energy ψ to maximise the invasion efficiency. Abdalrahman and Franz (2017) proposed an analytical model based on contact mechanics to describe the erythrocyte membrane deformation due to the ligand-receptor adhesion energy released during membrane indentation up to 10% of the merozoite length.

Mechanical markers, also referred to as mechanical phenotypes or mechanotypes, are emerging label-free markers that may offer the potential for use as drug targets (Qi et al., 2015). Here, mathematical and computational investigations can further clarify the mechanisms that determine the merozoite’s invasiveness and contribute to developing novel invasion blocking antimalarial compounds. One example, although demonstrated for viral entry into host cells, may be competitive ligand binding leading to receptor depletion and reduction adhesion energy for deformation of the cell membrane (Parveen et al., 2017).

As a first step, a detailed mathematical description of the biomechanical interactions between merozoite and erythrocyte during the invasion process is required, integrating the separate concepts of (i) the actomyosin motor force generation, (ii) spectrin network disruption and (iii) tight junction. Hence, this study aimed to develop an analytically model that describes the erythrocyte membrane mechanics during the entire merozoite entry and investigate with the model the effects of erythrocyte membrane deformation and phosphorylation on invasion forces and work.

## 2 Methods

The erythrocyte membrane wrapping mechanism was described analytically using the membrane theory of shells. (Note: Throughout the article, the term erythrocyte ‘membrane’ always refers to the lipid bilayer and the spectrin network, whereas some cases explicitly refer to the spectrin network only.)

A differential equation describing the equilibrium configuration of a circular erythrocyte membrane section and a rigid egg-shaped merozoite was solved numerically throughout the entire invasion process (Villaggio, 1997). The deformation of the erythrocyte membrane was described using merozoite shape parameters. Equilibrium and governing equations were formulated with deformation and constitutive law of the erythrocyte. The governing equations were numerically solved using the Runge Kutta fourth-order method to obtain stretches and stresses in the erythrocyte membrane.

The erythrocyte membrane wrapping mechanics was explored for two values of areal erythrocyte membrane strain, i.e. As ≤ 4% as generally accepted for quasi-static erythrocyte membrane deformation (Li et al., 2013) and As ≤ 51%. Additionally, the impact of the tight junction and merozoite-induced erythrocyte membrane damage on the invasion energetics were considered.

The following simplifying assumptions were used for the wrapping mechanics of a health erythrocyte during merozoite entry:

a. The erythrocyte membrane is homogeneous and mechanically isotropic.
b. The erythrocyte membrane, comprising lipid bilayer and spectrin network, has a small thickness of 0.01 μm, based on a thickness of the lipid bilayer alone of 4 to 5 nm (Baskurt and Meiselman, 2003; Hochmuth et al., 1983).
c. The region of the erythrocyte membrane involved in the wrapping is initially flat, considering a curvature of the erythrocyte membrane that is small compared to the merozoite surface at the attachment site.
d. The bending modulus of the erythrocyte membrane is small, and only the mid-plane strain due to loading modes other than bending is considered. This assumption is based on a very small ratio of membrane thickness to cell radius and the small ratio of bending to shear rigidity of less than 0.01 (Evans, 1983)
e. The merozoite is rigid, i.e., non-deformable.

The analytical model was implemented in MATLAB (The MathWorks, Inc, Natick, MA, USA) and the custom code files are available for download as described in the Data availability section.

### 2.1 Physical properties of an erythrocyte

A healthy mature human erythrocyte is a biconcave discoid shaped cell with a diameter of 7.8 μm and a thickness that varies from 0.8 to 2.6 μm. It is mainly composed of the cytoplasm surrounded by a membrane. In a healthy erythrocyte, the cytoplasm comprises a concentrated haemoglobin solution with a viscosity of approximately 6 to 10 mPa s (Dimitrakopoulos, 2012). Under normal physiological conditions, the haemoglobin viscosity has a negligible contribution to erythrocyte deformability, mainly determined by its membrane. However, at high viscosity of 650 mPa s (Mohandas and Chasis, 1993) the erythrocyte cytoplasm becomes a primary determinant of cellular deformability (Mohandas et al., 1979).

The erythrocyte membrane is a layered structure comprising the lipid bilayer and the membrane skeleton. The lipid bilayer is often considered a two-dimensional incompressible structure without shear resistance, whereas the underlying membrane skeleton is composed of a spectrin network that enables the erythrocyte membrane to resist shear loads (Dimitrakopoulos, 2012). The biconcave discoid shape of a healthy erythrocyte allows it to undergo large deformation of up to 250% of its original dimension while maintaining minimal areal membrane strain (Mohandas and Gallagher, 2008).

An erythrocyte has a volume of 94.5 μm^3^ and a surface area of 135.2 μm^2^ (Li et al., 2014). Compared to the surface area of a sphere with an identical volume, i.e. 98 μm^2^, the erythrocyte’s discoid shape increases its surface area by approximately 37 μm^2^, which is approximately 4.6 times the surface area of a merozoite of 8.0 μm^2^ (Dasgupta et al., 2014). This excess surface area suggests that the erythrocyte may wrap the merozoite entirely with a small areal strain in the membrane.

### 2.2 Merozoite geometry

The idealised ellipsoidal geometry of the merozoite is defined by:

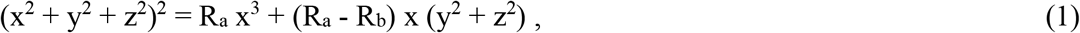

where R_a_ = 1 μm and R_b_ = 0.7 μm are shape parameters determining the size of the merozoite (Figure 1 b).

Using spherical coordinate parameterisation, the x, y, and z coordinates of the surface of the merozoite are expressed as (Dasgupta et al., 2014):

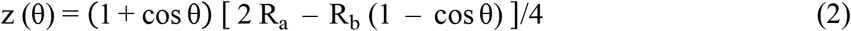

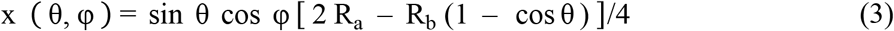

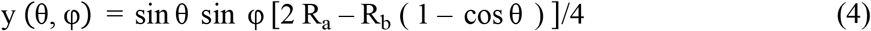

where θ and φ are polar angles.

The arc length on the egg-shaped merozoite surface is defined using the first fundamental form:

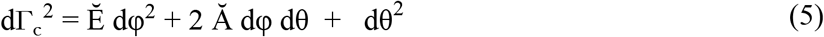

where Ě = r_φ_ × r_φ_, Ă = r_θ_ × r_φ_, Ğ = r_θ_ × r_θ_

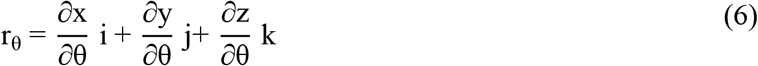

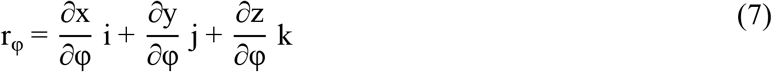

Letting ρ^2^ = (x^2^ + y^2^), the matrix of coefficients of the first form is defined:

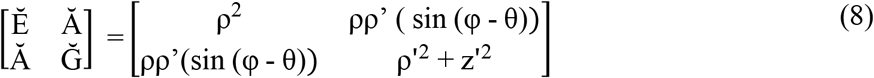

where a single prime represents the first partial derivative with respect to θ.

### 2.3 Deformation of erythrocyte membrane during merozoite wrapping

The erythrocyte membrane is assumed to be flat during the initial contact with the merozoite. During the wrapping of the merozoite, the main erythrocyte membrane strains and stresses are in the meridian and circumferential direction (Figure 1c), and the erythrocyte membrane deformation is parameterised using the merozoite’s shape parameters (Figure 1d). The shape parameters R_a_ and R_b_ can be used to investigate the impact of shape changes of the merozoite on its invasion efficiency.

The erythrocyte membrane deformations are described by meridian and circumferential stretch as follows:

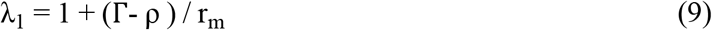

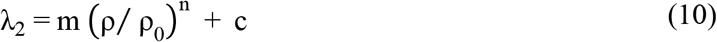

with

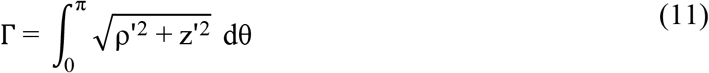

Here, ρ and z are the Cartesian coordinates of locations on the surface of the rigid merozoite; m, n, and c determine the magnitude of the principal erythrocyte membrane stretches, r_m_ is the radius of the circular erythrocyte membrane section, and Γ is the coverage length of the erythrocyte membrane on the merozoite surface in the meridian direction. (The subscripts 1 and 2 refer to the erythrocyte membrane’s meridian and circumferential principal directions, respectively, see Figure 1c.)

The meridian and circumferential deformation gradients are represented with the derivative of Eqns. (9) and (10):

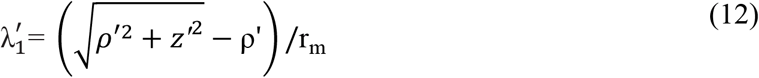

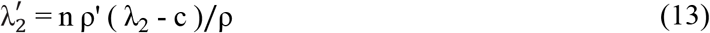

The local erythrocyte membrane bending is characterised by two principal curvatures, k_1_ and k_2_:

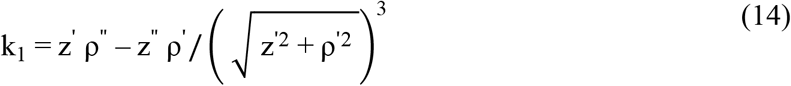

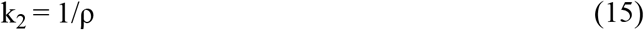

where the single and double prime represent the first and second partial derivatives with respect to θ.

### 2.4 Constitutive law of erythrocyte membrane

A typical erythrocyte model often comprises a Newtonian liquid cytoplasm surrounded by a thin membrane with a thickness of 0.01 μm (Hochmuth et al., 1973). The erythrocyte membrane is considered a mechanically isotropic, incompressible and hyperelastic solid (Villaggio, 1997).

The merozoite-induced erythrocyte membrane dissociation is represented by reducing the erythrocyte elastic modulus from 1 kPa to 0.5 kPa. The Mooney Rivlin strain energy density function per reference volume (Skalak et al., 1973) is:

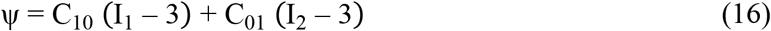

where C_10_ and C_01_ are empirical material parameters, I_1_ and I_2_ are the first and second strain invariants related to the principal stretch λ_1_ and λ_2_ by:

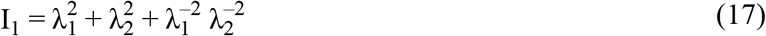

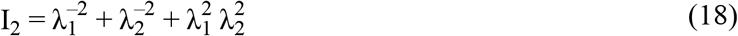

The material parameter C_10_ is related to the elastic modulus E of the erythrocyte membrane (Zhang and Zhang, 2011):

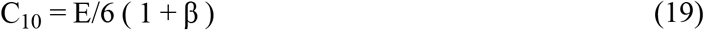

where β ranges from 0 to 0.5. The elastic modulus E is of the order 1 kPa (Hochmuth et al., 1973) and the ratio of two Mooney Rivlin material parameters is β = C_10_/C_01_ = 0.1 (Zhang and Zhang, 2011).

The erythrocyte membrane constitutive law is defined by:

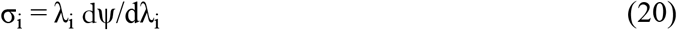

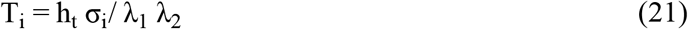

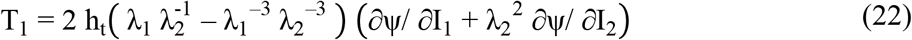

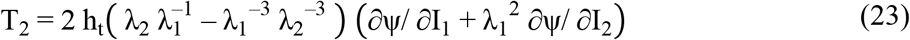

where h_t_ = 0.01 μm is the thickness of the erythrocyte membrane (Hochmuth et al., 1973).

The shear modulus G_s_ of the erythrocyte membrane is determined by:

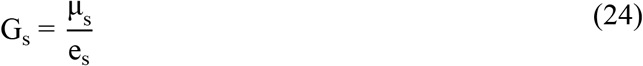

with

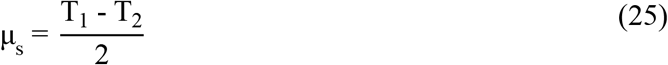

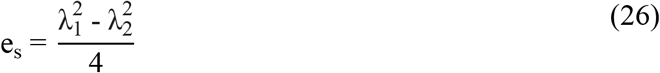

The areal strain A_s_ of the erythrocyte membrane is expressed by:

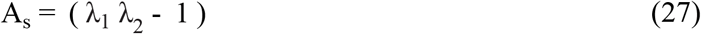

### 2.5 Equilibrium equations of erythrocyte membrane

For the contact region between the erythrocyte membrane and the rigid merozoite, the equilibrium equations are:

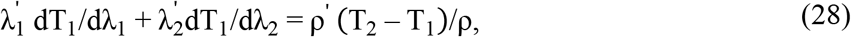

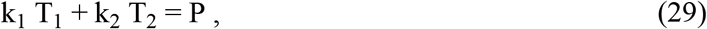

where P is the pressure exerted by the merozoite on the erythrocyte membrane surface that is directed normal to the erythrocyte membrane surface. The merozoite wrapping force F_w_ acting on the erythrocyte membrane surface and causing the erythrocyte membrane to wrap the merozoite is:

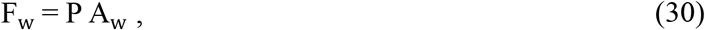

where A_w_ is the surface area of the merozoite wrapped by the erythrocyte membrane:

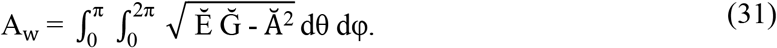

The total indentation force of the merozoite is:

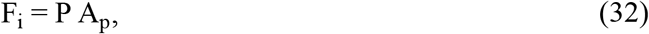

where A_p_ is the projected surface area of the erythrocyte membrane:

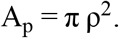

The indentation work, which is the energy required by the parasite to invade a human erythrocyte, is:

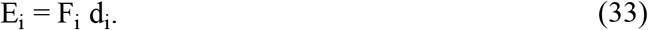

where d_i_ = (max(z) + z) is the indentation depth.

### 2.6 Governing equations of erythrocyte membrane

The governing differential equations for the contact region are:

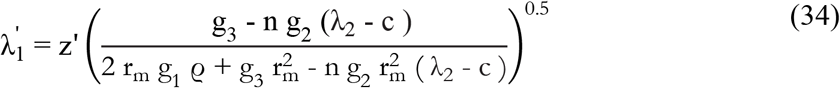

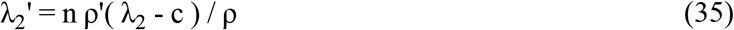

with

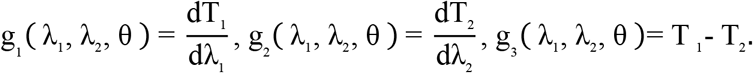

The coverage area of the erythrocyte membrane at each invasion step is:

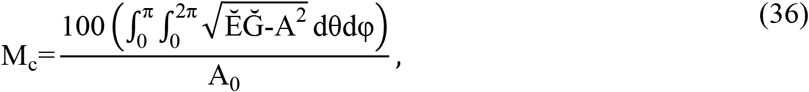

where A_0_ is the total surface area of the merozoite.

With the parameter values in Table 1, the solution of the governing equations can be obtained.

**Table 1:** Erythrocyte membrane parameters for simulating merozoite wrapping with small and large maximum areal membrane strain of A_s,max_ = 4% and 51%.

| $A_{s,\max}$<br>(%) | $E$<br>(kPa) | $C_{10}$<br>(Pa) | $C_{01}$<br>(Pa) | $r_m$<br>($\mu\text{m}$) | $c$ | $n$ | $\lambda_o$ |
| --- | --- | --- | --- | --- | --- | --- | --- |
| **4** | 1 | 183 | 18.3 | 50 | 1 | 1 | 1 |
| **51** | 1 | 183 | 18.3 | 4 | 1 | 1 | 1 |

### 2.7 Contribution of tight junction, erythrocyte membrane wrapping, and erythrocyte membrane damage to invasion energetics

The tight junction forms a mechanical link between the merozoite and erythrocyte membrane to facilitate membrane wrapping. It acts as an anchor location where the merozoite’s actomyosin motors pull the erythrocyte membrane. The tight junction is assumed to exert tension in the circumferential direction at any equilibrium point. In the erythrocyte membrane, a tension acting in the meridian direction is assumed to resist the forces applied by the merozoite. The tensions in the meridian direction T1 and circumferential direction T2 are used to describe the energetics of the tight junction and the erythrocyte membrane. The parameter values in Table 1 consider the contributions of the tight junction and the erythrocyte membrane wrapping during small and large strain deformation of the erythrocyte membrane.

After attaching to the erythrocyte, the merozoite induces phosphorylation that locally dissolves the spectrin network, forming the erythrocyte membrane’s skeleton. This process changes the erythrocyte membrane’s mechanical properties (Zuccala and Baum, 2011). For example, phosphorylation of band 4.1 weakens the connections between band 4.1, spectrin and actin, which leads to membrane destabilisation (Betz et al., 2009). The phosphorylation of β spectrin also leads to decreased membrane stability (Manno et al., 1995); tyrosine phosphorylation of the SLC4A1 protein weakens the attachment of SLC4A1 to the underlying spectrin network and increases the membrane mobility by altering ankyrin binding (Ferru et al., 2011). To assess the impact of membrane damage, the erythrocyte membrane elastic modulus was reduced from 1 kPa to 0.5 kPa.

## 3 Results

The developed model was applied to simulate the wrapping by an erythrocyte membrane. Two cases of maximum erythrocyte membrane areal strain were considered, i.e. (i) A_s,max_ = 4% as a generally accepted upper threshold for initiation of cell lysis during quasi-static deformation of the membrane (Li et al., 2013), and (ii) A_s,max_ = 51% to investigate larger deformations with an upper threshold of 40% yield areal strain of the erythrocyte membrane (Li et al., 2013). The effect of local membrane damage due to phosphorylation by the merozoite was considered by reducing the erythrocyte membrane’s elastic modulus from 1 kPa to 0.5 kPa.

The maximum areal strain in the erythrocyte membrane during merozoite wrapping was implemented with a large circular membrane section with radius r_m_ = 50 μm for A_s,max_ = 4% and a small membrane section with r_m_ = 4 μm for A_s,max_ = 51%, along with the parameter values specified in Table 1.

The merozoite wrapping with A_s,max_ = 4% induced maximum principal stretches of λ_1,max_ = 1.04 in the meridian direction and λ_2,max_ = 1.000046 in circumferential direction of the erythrocyte membrane (Figure 2). The maximum circumferential stretch λ_2,max_ occurred at a meridian stretch of λ_1_ = 1.023 (Figure 2 a). Permitting a higher areal membrane strain of A_s,max_ = 51% during merozoite wrapping resulted in a substantially increased maximum principal membrane stretches of λ_1,max_ = 1.5 and λ_2,max_ = 1.0375, the latter occurring at λ_1_ = 1.26 (Figure 2 b). The circumferential and meridian stretch in the erythrocyte membrane during the invasion process is illustrated in (Figure 2 c-f).

**Figure 2:**
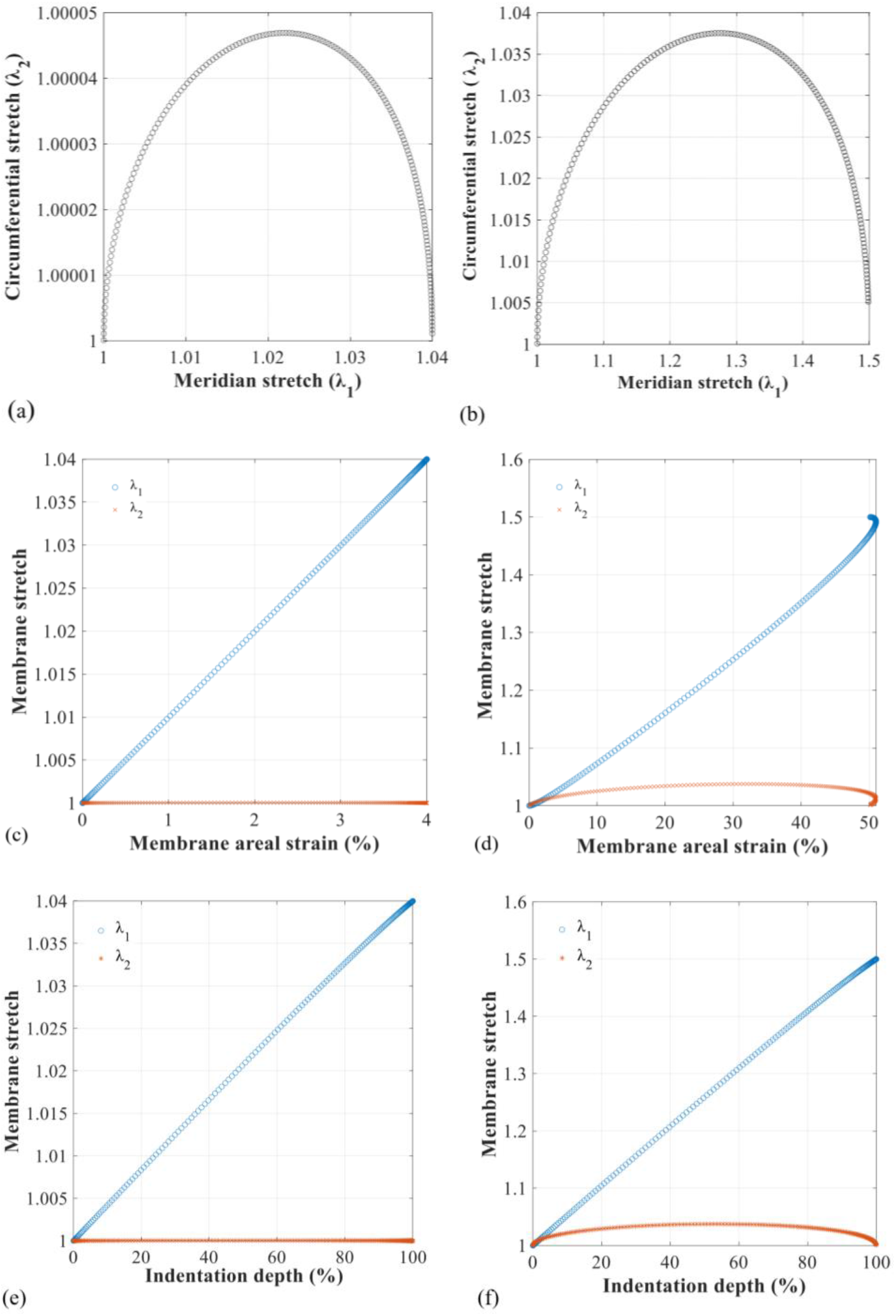
Numerical solution of principal stretches in the erythrocyte membrane for the complete wrapping of the merozoite. Circumferential stretch λ_2_ versus meridian stretch λ_1_ for (a) small maximum areal strain of A_s,max_ = 4% and (b) large maximum areal strain of A_s,max_ = 51%. The meridian stretch λ_1_ and circumferential stretch λ_2_ versus areal strain in the erythrocyte membrane for A_s,max_ = 4% (c) and 51% (d). The meridian stretch λ_1_ and circumferential stretch λ_2_ versus indentation depth normalised to merozoite length L_m_ for A_s,max_ = 4% (e) and 51% (f).

The tension induced by the circumferential and meridian stretches for small areal strain up to A_s,max_ = 4% reflects a near-linear constitutive response of the erythrocyte membrane (Figure 3 a). For large areal strain, A_s,max_ = 51%, the erythrocyte membrane’s nonlinear elastic response with strain softening is apparent (Figure 3 b).

**Figure 3:**
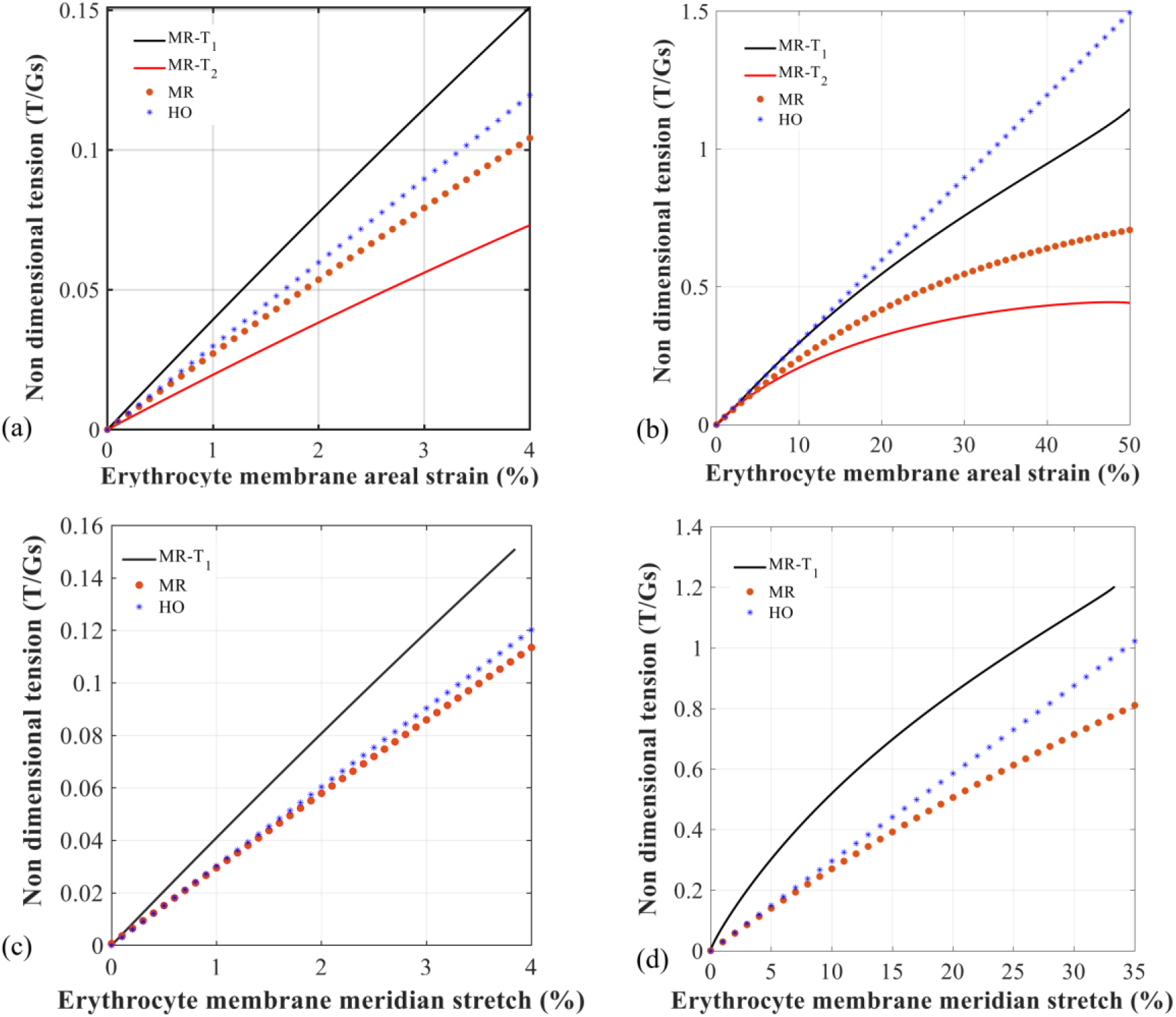
Non-dimensional isotropic principal tension versus areal strain (a, b) and meridian strain (c, d) in the erythrocyte membrane during merozoite wrapping for the maximum areal strain of A_s,max_ = 4% (a, c) and 51% (b, d). Solutions for the meridian (MR-T1) and circumferential tension (MR-T2) obtained with the developed Mooney-Rivlin-based erythrocyte membrane constitutive model (Eqns. (22) and (23)) compared to solutions of non-dimensional tension based on the linear elastic Hooke’s law (HO) and the nonlinear elastic Mooney-Rivlin law (MR) from Omori et al. (2011).

The model’s prediction of the non-dimensional meridian membrane tension T1 agrees better with Hooke’s law non-dimensional tension than the Mooney-Rivlin-based non-dimensional tension for small membrance deformation with A_s,max_ = 4% (Omori et al., 2011) (Figure 3 a, c), and for larger membrane strain with A_s,max_ = 51 (Figure 3 b, d).

The erythrocyte membrane’s strain softening was also apparent by a decreasing shear modulus with increasing areal strain (Figure 4 a). The meridian and circumferential membrane stresses associated with an areal strain of A_s,max_ = 4% were σ_1_ = 63.1 Pa and σ_2_ = 30.5 Pa (see Figure 4 b).

**Figure 4:**
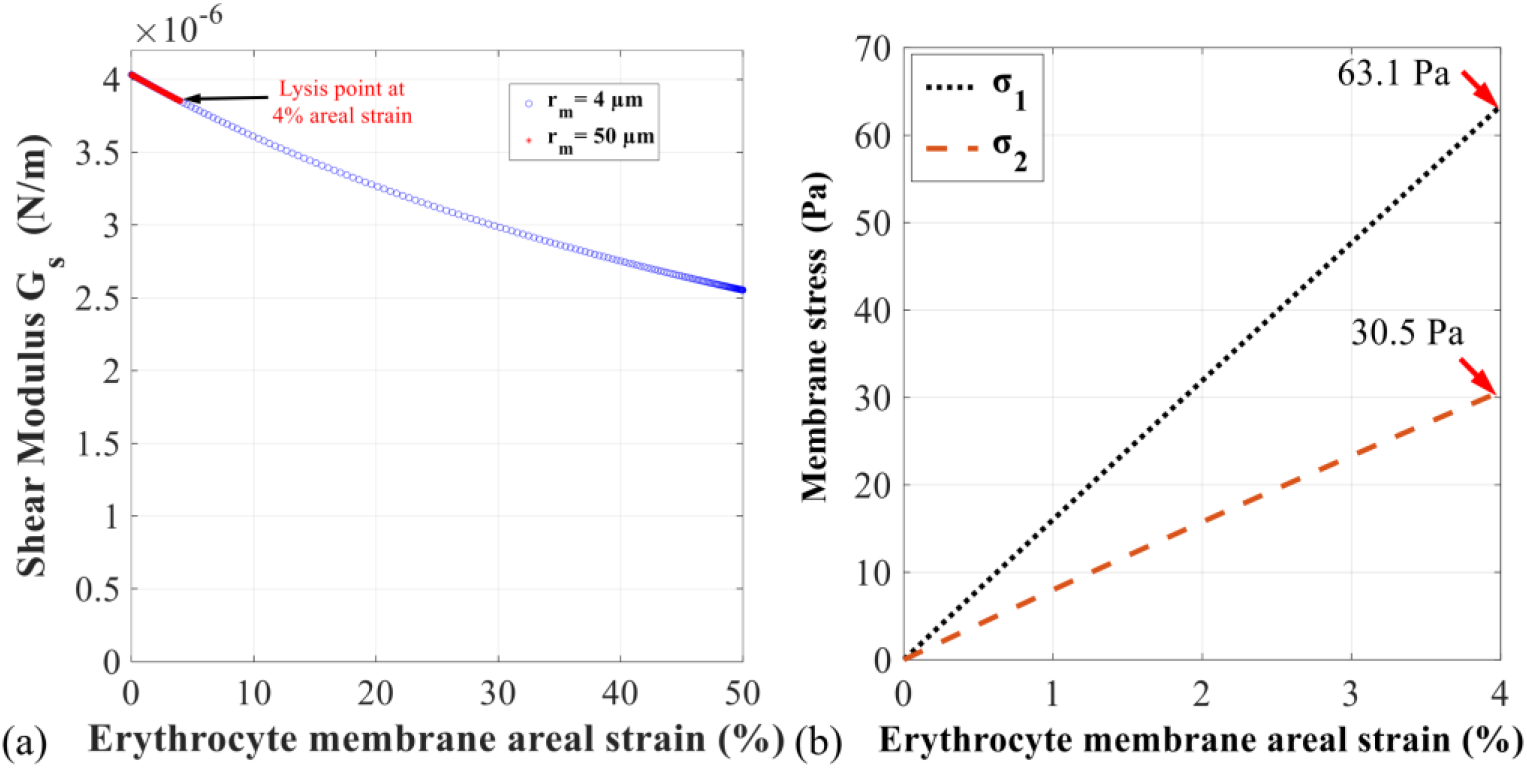
(a) Shear modulus versus areal strain of the erythrocyte membrane during complete wrapping of the merozoite. The decrease of the shear modulus with increasing areal strain represents strain softening. (b) Meridian and circumferential stress versus areal strain in the erythrocyte membrane during complete merozoite wrapping. The stress associated with the assumed lysis threshold of the areal strain of A_s,max_ = 4% is σ_1_ = 63.1 Pa in the meridian and σ_2_ = 30.5 Pa in the circumferential direction.

The total indentation force F_i_ during membrane wrapping was predicted to increase nearly linearly up to a normalised indentation depth of 85.2% marking the position of the merozoite’s maximum diameter and decrease after that. The indentation force was higher for the large (A_s,max_ = 51%) than the small membrane strain (A_s,max_ = 4%); however, the F_i_ maximum occurred at the same indentation depth (see Figure 5 a). The total wrapping force was predicted to increase exponential during the late stages of the invasion process. These results illustrate that a large surface area of the erythrocyte membrane involved in the wrapping leads to smaller magnitudes of total wrapping and indentation forces.

**Figure 5.**
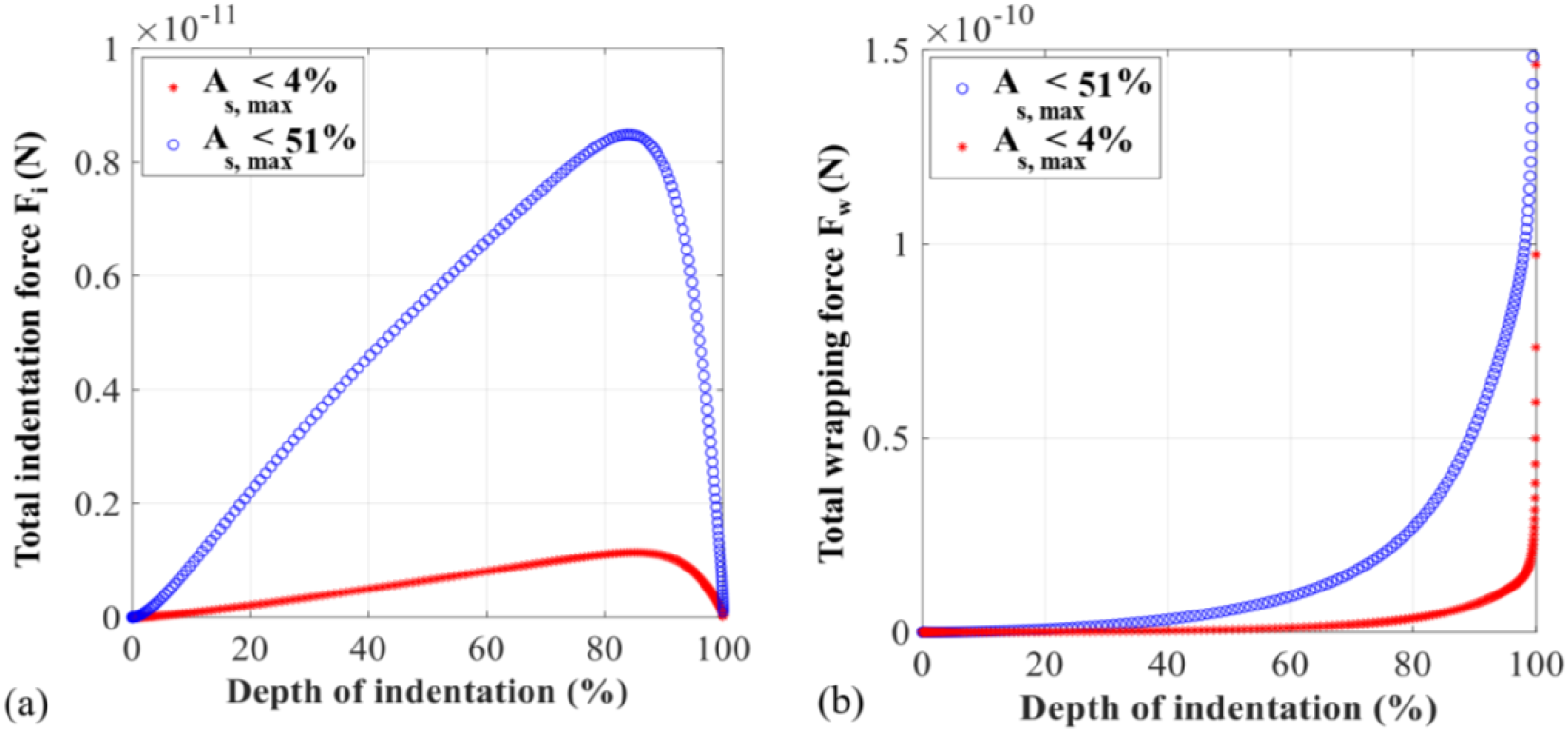
Forces involved in merozoite wrapping versus normalised indentation depth d_i_/L_m_ for small (A_s,max_ = 4%) and large maximum areal strain (A_s,max_ = 51%) in the erythrocyte membrane. The normalised indentation depth d_i_/L_m_ relates the indentation depth d_i_ to the merozoite length L_m_, where d_i_/L_m_ = 100% corresponds to d_i_ = L_m_ and full indentation of the merozoite (see also Figure 1d). (a) Total indentation force F_I_ acting in the direction of the merozoite’s long axis. (b) Total wrapping force F_W_ acting normal to the surface of the merozoite.

The total indentation work E_i_ required by the merozoite to overcome the barrier energy of the erythrocyte membrane was predicted to increase with increasing indentation depth. E_i_ reaches a maximum of E_i_ = 1.92×10^-18^ J for A_s,max_ = 4% and E_i_ = 1.40×10^-17^ J for A_s,max_ = 51% at the same normalised indentation depth as the total indentation force, i.e. at 85.2% (Figure 6). A reduction of the erythrocyte membrane’s elastic modulus from 1 to 0.5 kPa to simulate weakening due to phosphorylation of the spectrin network led to a reduction of the total indentation work by 50% to E_i_ = 0.97×10^-18^ J for A_s,max_ = 4% and E_i_ = 0.70×10^-17^ J for A_s,max_ = 51% (Figure 6).

**Figure 6:**
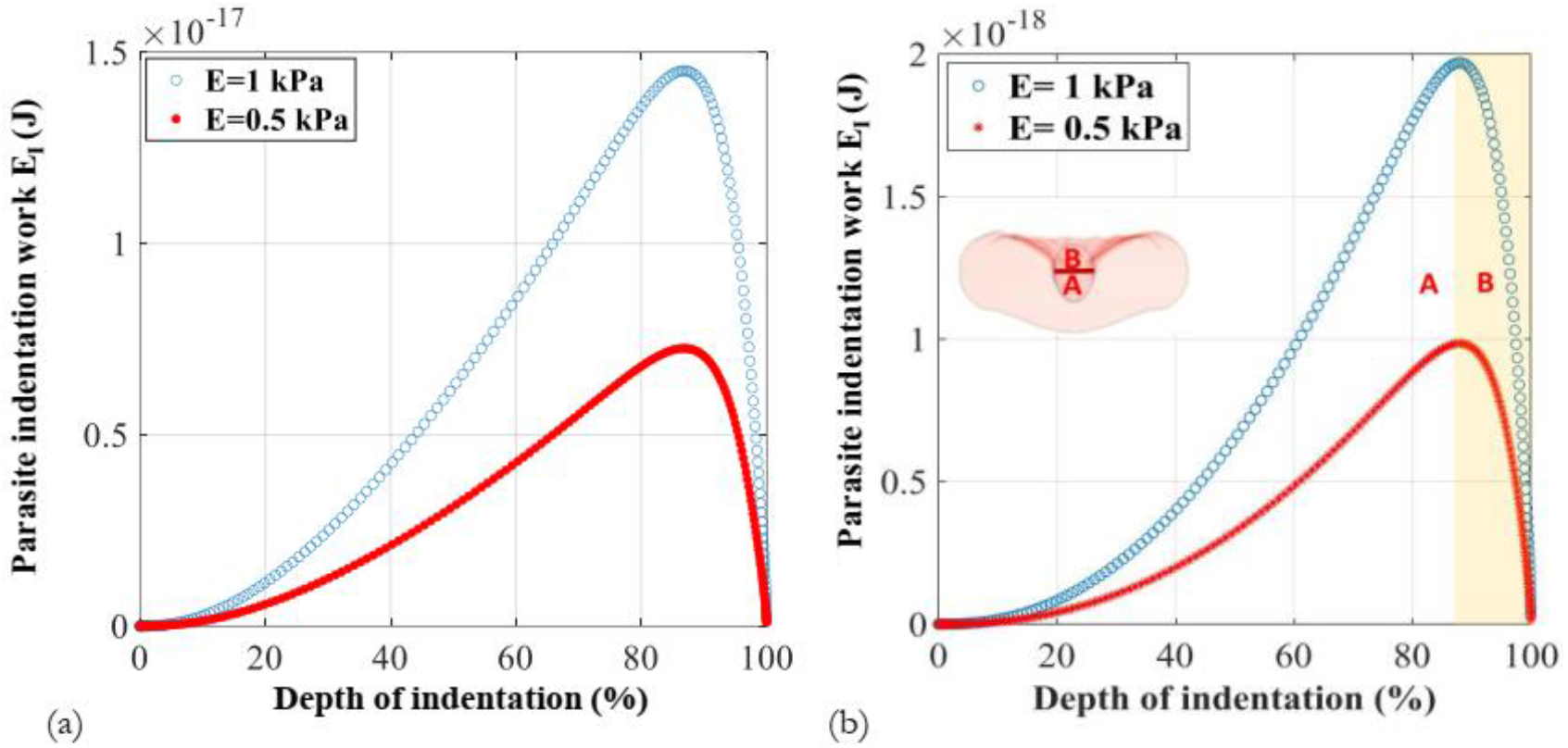
Indentation work E_i_ required for merozoite wrapping versus normalised indentation depth d_i_/L_M_ without and with phosphorylation of the spectrin network (represented by reducing the membrane’s elastic modulus from 1 kPa to 0.5 kPa) for A_s,max_ = 51% (a) and 4 % (b). Phosphorylation decreased E_i_ by 50%. In (b), labels A and B represent the indentation depth regions for which the merozoite and tight junction, respectively, provide the indentation energy, with the change occurring at a normalised indentation depth of 85.2%.

## 4 Discussion

An analytical model for wrapping a malaria merozoite by an erythrocyte was developed. The effects of the erythrocyte membrane area available for wrapping, membrane damage through phosphorylation of the spectrin network, and the tight junction between merozoite and erythrocyte membrane in the invasion mechanics were investigated. The tight junction was defined by a line tension acting in the circumferential direction of the erythrocyte membrane with an initial radius ρ_o_. The total energy required for a successful invasion, i.e. the internalisation of the parasite, was determined by computing the total work done by the merozoite, representing the contributions of the tight junction and the erythrocyte membrane.

The results of the developed analytical model indicate that during the early stage of active invasion, the work done by the merozoite increases progressively and is stored as internal energy in the tight junction and the erythrocyte membrane. This energy is utilised during the late phase of the invasion, facilitating the sealing of the erythrocyte membrane during the late invasion stage.

Two erythrocyte membrane wrapping cases were assessed, distinguished by the maximum areal strain permitted in the erythrocyte membrane during the invasion, namely A_s,max_ = 4% and 51%. The strain limits were implemented with circular erythrocyte membrane areas of different sizes involved in the wrapping; a large area with r_m_ = 50 μm for the small strain limit and a small area with r_m_ = 4 μm for the large strain limit. it was observed in the predictions that the merozoite wrapping force F_w_ and the total indentation force F_i_ were lower for low areal strain in the erythrocyte membrane (and large wrapping membrane area) than for the large strain (and small membrane area) (Figure 5). This finding suggests that the merozoite’s invasiveness efficiency depends on the excess surface area of the erythrocyte membrane available due the erythrocyte’s biconcave shape and the areal strain induced in the membrane during the wrapping process.

It has been reported that discoid-shaped erythrocytes are more deformable than ellipsoidal or spherical erythrocytes, often found with hereditary disorders such as elliptocytosis and spherocytosis (Li et al., 2018). The increased deformability stems from the discoid’s larger surface-to-volume ratio (S/V) than the spherical and elliptic shapes (Tomaiuolo, 2014). However, it has been unclear whether and how the different erythrocyte shapes affect the invasion of merozoites. Predictions with the developed model revealed that the required invasion force and energy are lower for a large erythrocyte membrane area than for a small membrane area involved in the wrapping interaction (Figure 5). The lower invasion force and energy may promote the merozoite invasion and increase invasion efficiency. This finding can also partly explain why individuals with abnormal erythrocyte shapes, e.g., caused by spherocytosis, elliptocytosis, and sickle cell anaemia, are less susceptible to malaria. However, further analysis is needed to quantify the impact of surface-to-volume ratio on the merozoite invasiveness.

The effect of merozoite-induced phosphorylation of the erythrocyte’s spectrin network and membrane damage in the invasion mechanics was assessed by reducing the elastic modulus of the erythrocyte membrane in the Mooney Rivlin law from 1 kPa for the intact membrane to 0.5 kPa for the membrane with damage. This change led to a decrease in the required invasion work and energy (Figure 6), suggesting that the phosphorylation of the erythrocyte membrane supports the merozoite invasion and increases invasion efficiency. Varying the reduced membrane elastic modulus due to phosphorylation in future studies may provide information on changes in the invagination mechanics with increasing phosphorylation damage to the erythrocyte membrane during merozoite entry.

The mechanical erythrocyte membrane response in the meridian direction predicted by the developed model agrees better with the solution based on Hooke’s law than the original Mooney-Rivlin model (Omori et al., 2011). The noticeable differences between the developed model’s predictions and the previous numerical data are due to the different loading modes, i.e. merozoite-induced stretch for the developed model versus biaxial stretch for the Hooke’s law and Mooney-Rivlin solutions by Omori et al. (2011) (Figure 3).

A recent study indicated that the wrapping energy for the control population is in the range of 1.40 ×10-^17^ J (Introini et al., 2022), which is equal to the parasite indentation work predicted by the developed analytical model for a 51% areal membrane strain; hence supporting the validity of the developed model.

The developed analytical model has some limitations, i.e. the restriction to 2D analysis and the absence of the erythrocyte cytoplasm, and the extension of the model is recommended for future work. For demonstration of the model in the current study, the stiffness reduction due to phosphorylation was applied in the entire erythrocyte membrane sections modelled. However, since phosphorylation takes place very localised, the assumption of uniform stiffness reduction in the membrane holds more for the small than the large membrane section. For implementation and use of the analytical model in future studies, the area of phosphorylation can and should be adjusted to represent the local effect of phosphorylation. In this case, the assumption of uniform stiffness reduction is not applicable and required, and the membrane heterogeneity can be considered in the model.

## 5 Conclusions

The developed analytic model is the first capable of predicting the invagination forces and energetics generated by the merozoite actomyosin machinery and erythrocyte membrane and assessing the merozoite’s invasiveness based on the erythrocyte membrane’s shape and mechanics. The current study demonstrated the model’s feasibility for providing new knowledge on the physical mechanisms of the merozoite invasion process that contribute to the invasion efficiency. While the current model focuses on the plasmodium falciparum merozoite, it can be easily applied to other malaria parasite variants by changing the shape parameters of the merozoite. As such, the model may be beneficial in the discovery of new invasion-blocking anti-malaria drugs, especially following extension in the future. The implementation of the analytical model with an enhanced model for erythrocyte membrane damage due to phosphorylation into a finite element model of an erythrocyte would be one example. Such models may facilitate predictive *in silico* studies to develop more efficient assays for anti-malaria drugs that are based on quantifying changes in erythrocyte mechanical properties induced by localised phosphorylation erythrocyte membrane damage.

## Data availability

Custom MATLAB code files of the model and numerical data supporting the results presented in this article are available on the University of Cape Town’s institutional data repository (ZivaHub) under https://doi.org/10.25375/uct.19641306 as Msosa C, Abdalrahman T, Franz T. Data for an analytical model describing the mechanics of erythrocyte membrane wrapping during active invasion of a plasmodium falciparum merozoite, Cape Town, ZivaHub, 2022, DOI 10.25375/uct.19641306.

## Nomenclature

A_0_: Undeformed erythrocyte membrane area
A_p_: Projected erythrocyte membrane area
A_s_: Areal strain of the erythrocyte membrane
A_s,max_: Maximum areal strain of the erythrocyte membrane
A_w_: Wrapped surface area of the merozoite
Ă: 3D merozoite shape parameter
c: Stretch parameter
C_01_: Material parameter
C_10_: Material parameter
d_i_: Indentation depth
E: Elastic modulus
E_i_: Parasite indentation work
Ě: 3D merozoite shape parameter
e_s_: Shear strain
F_i_: Parasite indentation force
F_w_: Parasite wrapping force
G_s_: Shear modulus
Ğ: 3D merozoite shape parameter
h_t_: Erythrocyte membrane thickness
I_1_: First strain invariant
I_2_: Second strain invariant
k_1_: Meridian curvature
k_2_: Circumferential curvature
m: Stretch parameter
M_c_: Coverage area of the erythrocyte membrane
MR-T_1_: Meridian tension obtained using the Mooney Rivlin law
MR-T_2_: Circumferential tension obtained using the Mooney Rivlin law
n: Stretch parameter
P: Pressure acting on the erythrocyte surface
R_a_: Merozoite shape parameter
R_b_: Merozoite shape parameter
r_m_: Radius of erythrocyte membrane section
T_1_: Principal meridian tension
T_2_: Principal circumferential tension
x: x coordinate of a point on the merozoite surface
y: y coordinate of a point on the merozoite surface
z: z coordinate of a point on the merozoite surface
β: Ratio of material parameters
Γ: Linear coverage of the erythrocyte membrane
Γ_c_: Arch length on the merozoite surface
θ: Polar angle defining the merozoite shape
λ_1_: Principal meridian stretch
λ_1,max_: Maximum principal meridian stretch
λ_2_: Principal circumferential stretch
λ_2,max_: Maximum principal circumferential stretch
μ: Dynamic viscosity
μ_0_: Initial shear modulus
μ_s_: Shear stress
v: Poisson’s ratio
ρ: Polar coordinate point on the merozoite surface
ρ_0_: Initial polar coordinate of a point on the merozoite surface at which the tight junction forms
σ_1_: Principal meridian stress
σ_2_: Principal circumferential stress
ψ: Mooney Rivlin strain energy density function per reference volume
φ: Polar angle defining the merozoite shape

